# Multiple Sites on SARS-CoV-2 Spike Protein are Susceptible to Proteolysis by Cathepsins B, K, L, S, and V

**DOI:** 10.1101/2021.02.17.431617

**Authors:** Keval Bollavaram, Tiffanie H. Leeman, Maggie W. Lee, Akhil Kulkarni, Sophia G. Upshaw, Jiabei Yang, Hannah Song, Manu O. Platt

**Affiliations:** Wallace H. Coulter Department of Biomedical Engineering at Georgia Institute of Technology & Emory University; Peking University

**Author notes:** To whom correspondence should be addressed: Manu O. Platt, Ph.D., 950 Atlantic Drive, Suite 3015, Atlanta GA, 30332.

**Keywords:** Molecular docking, cathepsin, COVID-19, proteolysis, inflammation, viral entry, extracellular matrix remodeling, computational modeling

## Abstract

SARS-CoV-2 is the coronavirus responsible for the COVID-19 pandemic. Proteases are central to the infection process of SARS-CoV-2. Cleavage of the spike protein on the virus’s capsid causes the conformational change that leads to membrane fusion and viral entry into the target cell. Since inhibition of one protease, even the dominant protease like TMPRSS2, may not be sufficient to block SARS-CoV-2 entry into cells, other proteases that may play an activating role and hydrolyze the spike protein must be identified. We identified amino acid sequences in all regions of spike protein, including the S1/S2 region critical for activation and viral entry, that are susceptible to cleavage by furin and cathepsins B, K, L, S, and V using PACMANS, a computational platform that identifies and ranks preferred sites of proteolytic cleavage on substrates, and verified with molecular docking analysis and immunoblotting to determine if binding of these proteases can occur on the spike protein that were identified as possible cleavage sites. Together, this study highlights cathepsins B, K, L, S, and V for consideration in SARS-CoV-2 infection and presents methodologies by which other proteases can be screened to determine a role in viral entry. This highlights additional proteases to be considered in COVID-19 studies, particularly regarding exacerbated damage in inflammatory preconditions where these proteases are generally upregulated.

## Introduction

Proteases are central to the infection process of SARS-CoV-2^1^, the coronavirus responsible for the COVID-19 pandemic^2^. Cleavage of the spike protein on the virus’s capsid into S1/S2 subunits causes the conformational change that leads to membrane fusion and viral entry into the target cell^3^. This can happen at the cell surface through the action of the membrane bound protease, TMPRSS2^4^ and soluble, extracellular furin^5^, and also after endocytosis of the virion facilitating endolysosomal escape through proteolytic activity of cysteine cathepsins B and L^6^ in acidic microenvironments^1^. These proteases have been the primary focus and implicated in SARS-CoV or SARS-CoV-2 infection^7^, but a number of other proteases may be contributing to spike protein activation, but have not yet been implicated^8^. Since inhibition of one protease may not be sufficient to block SARS-CoV-2 entry into cells^4^, other proteases that may play an activating role and hydrolyze the spike protein must be identified.

Cathepsins B and L have already been implicated in cleavage of SARS-CoV-2 spike protein^4^, and there have been strong suggestions to prioritize cathepsin L inhibitors for COVID-19 therapies that have already been developed — some are even FDA approved^9^. Cathepsins K, S, and V share 60% sequence homology with cathepsin L with cathepsin V actually sharing 80% sequence homology with cathepsin L^10^. These cathepsins are also promiscuous with their substrates, suggesting cross-reactivity^11^. Here, we test the hypothesis that the spike protein might have sites susceptible to cleavage by cathepsins K, S, or V and suggest additional mechanisms by which proteases are involved in SARS-CoV-2 infection.

SARS-CoV-2 infection and severity of COVID-19 were shown to be elevated in patients with pre-existing conditions, resulting in a greater risk of mortality in those patients^12^. These pre-existing conditions include diseases that are associated with upregulation of cathepsins K, L, S, and V such as diabetes^13–15^, hypertension^16–18^, sickle cell disease^19,20^, cardiovascular disease^14,21^, and emphysema^22,23^. Together, this led us to investigate spike protein amino acid sequences preferred in the active site of these cathepsins for peptide bond hydrolysis.

Here, we used an unbiased approached, applying computational methods to determine the potential of other closely related cysteine cathepsins to bind to and cleave the spike protein. Computational analyses of the SARS-CoV-2 genome-encoded protease necessary for viral replication^24^ suggested inhibitor molecules and strategies, but the focus here was to investigate human proteases that could also be targeted for inhibition. We used a program we previously developed called PACMANS, **P**rotease-**A**se **C**leavages from **M**EROPS **AN**alyzed **S**pecificities^25^ to determine putative cleavage site locations on SARS-CoV-2 spike protein. This algorithm works with users’ input of a substrate peptide’s amino acid sequence for cleavage by an identified protease that is included in the MEROPS database^26^. PACMANS uses a sliding-window approach, where individual sub-sequences that fill the active site pocket are scored, based on amino acids preferred at the active site of the particular protease for maximal hydrolytic ability against a region on the input substrate to generate scores for preferred cleavage sites. This pocket slides along the length of the substrate amino acid sequence, scores all possible 8-amino acid sub-sequences, then ranks them by the normalized version of this score. PACMANS also calculates a normalized score to enable comparisons across multiple proteases on the same substrate. Coupling sequence analysis with molecular docking tools, together, are an unbiased approach presented here to rationally identify other protease targets for new therapeutics against SARS-CoV-2, its infectivity, disease severity, and potentially longer-term complications^27^.

## Results

### Bioinformatic analyses using the protein sequence of spike protein indicates susceptibility to multiple cysteine cathepsins beyond those previously identified

Using PACMANS, possible putative cleavage sites on SARS-CoV-2 spike protein, the substrate, were identified with furin and cathepsins B, K, L, S, and V as the proteases. Furin, and cathepsins B and L have already been identified to cleave spike protein in previous studies^4,5^. From PACMANS analysis, furin had the highest scoring cleavage site and identified its cleavage at the RRAR / SVAS site known to be the activating cleavage at S1/S2 site that causes conformational change and membrane fusion^5^ (Table 1). This confirms that PACMANS does identify valid cleavage sites along the full length of the spike protein based solely on the amino acid sequence and its susceptibility to hydrolysis by these proteases. Cathepsin B, like furin, also prefers basic amino acid residues at its active site, and scored high at multiple sites on spike protein. Putative cleavage sites for cathepsins B, K, L, S, and V are included in the rank ordered list sorted by their normalized scores. Cleavage sites were identified in multiple domains of the spike protein (Table 1).

**Table 1.**
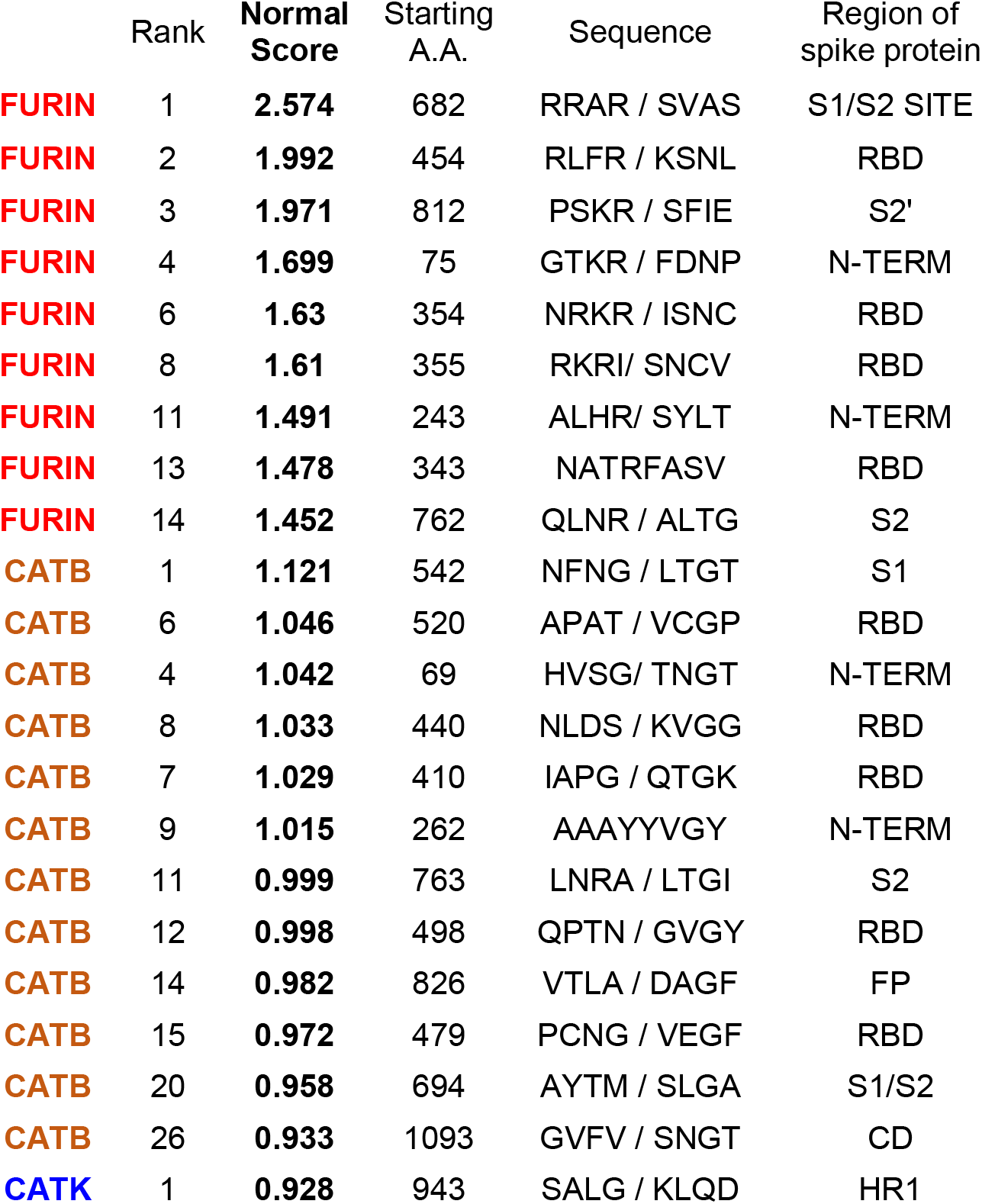
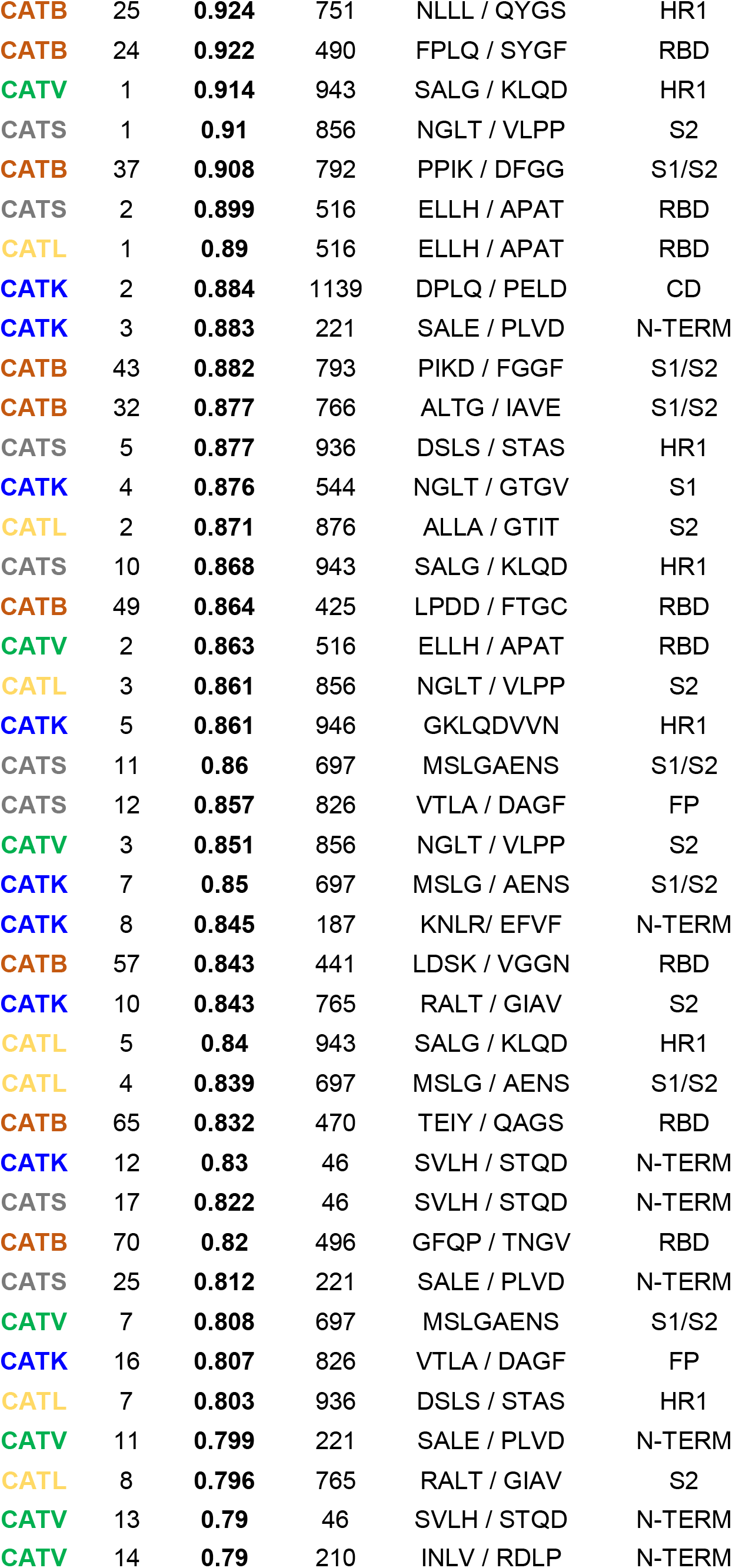
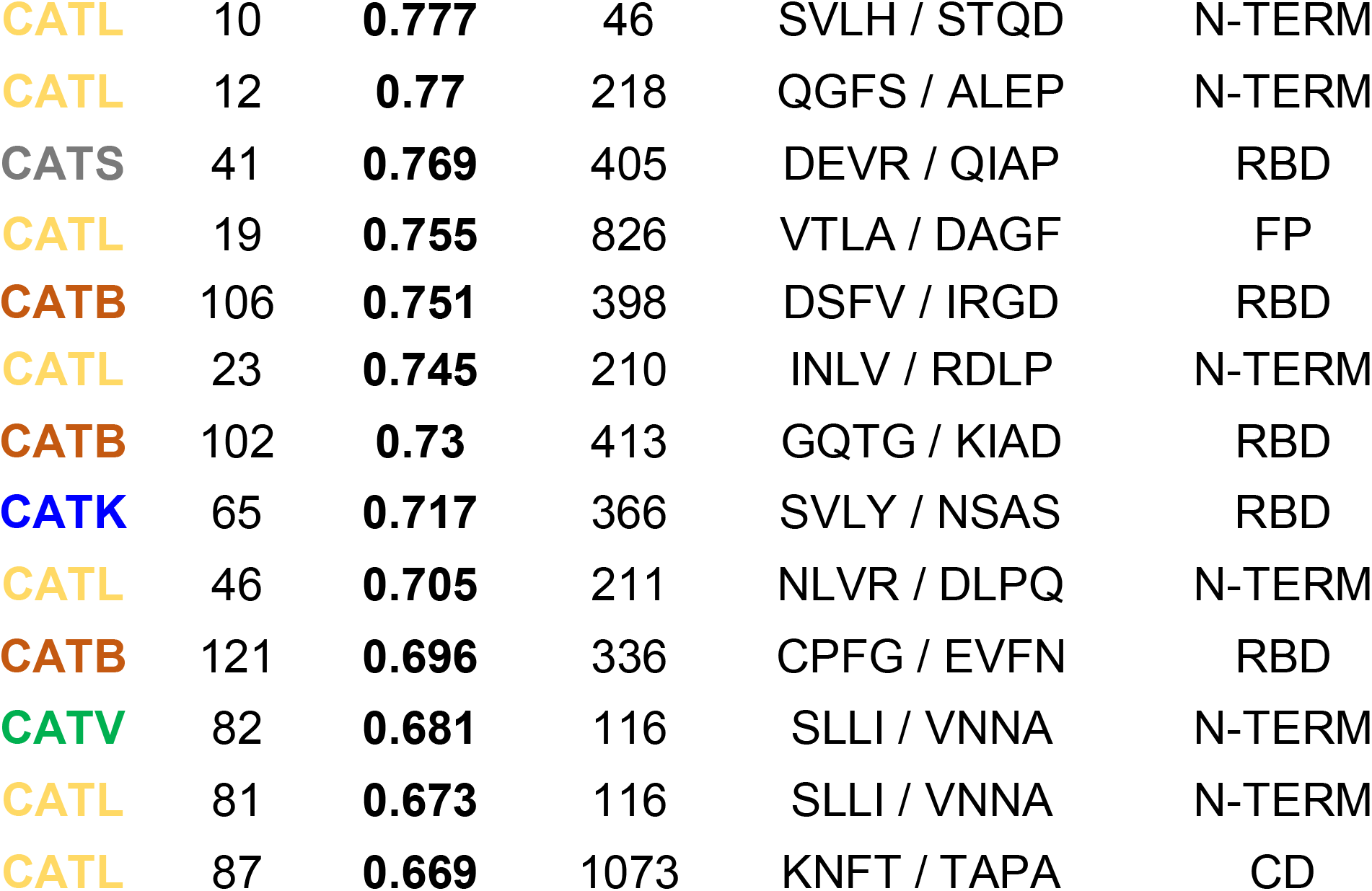
Results of PACMANS and docking analysis to identify preferred cleavage sites by multiple proteases on SARS-CoV-2 spike protein. Sequences for SARS-CoV-2 spike protein cleavage sites by six proteases were identified by PACMANS and sorted by their PACMANS normalized score. Ranks are for each individual protease on spike protein, based on their normalized score and likelihood of cleavage (lower the rank number, higher likelihood of cleavage). The ‘/’ indicates the hydrolytic site with amino acids on either side. Based on amino acid number, the region of the spike protein is also identified. Receptor binding domain (RBD), N-terminal domain (N-Term), Fusion peptide (FP), connector domain (CD), and heptad repeat (HR).

High ranking cleavage sites in the S1/S2 region was also identified for cathepsin K at amino acid 700 at the S2’ site which was the 7 highest rank according to PACMANS. A number of sequences were identified as highly ranked for their susceptibility to multiple proteases, indicating a strong likelihood of cleavage by one or more proteases. Ranks indicate the preferred site on spike protein for each specific protease, so each protease has a rank#1 for highest preference in the table for cleaving spike protein. The normalized score allows for comparison across all the proteases for the same substrate. There were potential cleavage sites in all regions of the spike protein including the receptor binding domain, the S1/S2 cleavage domain for spike protein activation, N-terminus, C-terminus, and heptad repeat domains by cathepsins B, K, S, V and L. Cathepsin B’s highest score region was in the S1 domain. Cathepsins K and V had the same top ranked site in the heptad repeat region, cathepsin S was highest scored in the S2 region, and cathepsin L was in the receptor binding domain.

### Molecular docking analyses corroborate protease active site binding at identified cleavage sites

Since PACMANS only analyzes the sequence of amino acids, consideration for the three-dimensional structure was helpful to determine access of the protease active sites to dock within close proximity of the spike protein sequence for cleavage is necessary. To assess this separately in an unbiased manner, furin, and cathepsins B, K, L, S, and V were used in molecular docking studies in ClusPro 2.0 to identify spike protein locations where the proteases’ active site would be sufficiently close to the SARS-CoV-2 spike protein to bind. Priority was given to those models with the protease catalytic triad facing the SARS-CoV-2 spike protein with distances near 10 Å or less. Both open and closed conformations of the spike protein were used in the analysis. From the open conformation, molecular docking sites were compiled onto one spike protein trimer and shown from multiple viewpoints in figure 1A - C. Cathepsins L and K had the closest binding distances with cathepsin L being 3.6 Å from V213 (N-terminal domain) and 5.8 Å from A123 (N-terminal domain), and cathepsin K being 3.8 Å from A372 (receptor binding domain). Cathepsin B was 10 Å from E340 (receptor binding domain). Cathepsin V’s closest distance was 12.2 Å from R214 (N-terminal domain) and 14.9 Å from A123 (N-terminal domain). From the closed conformation (Fig 1 D-F), again, cathepsins K and L were the closest distances with cathepsin K only 6.2 Å from V367 (receptor binding domain), and cathepsin L being 5.1 Å from R214 (N-terminal domain) and 5.3 Å from A123 (N-terminal domain). Cathepsin V again was less close, but still produced calculated distances of 11 Å from R214 (N-terminal domain) and 10 Å from N122 (N-terminal domain).

**Figure 1.**
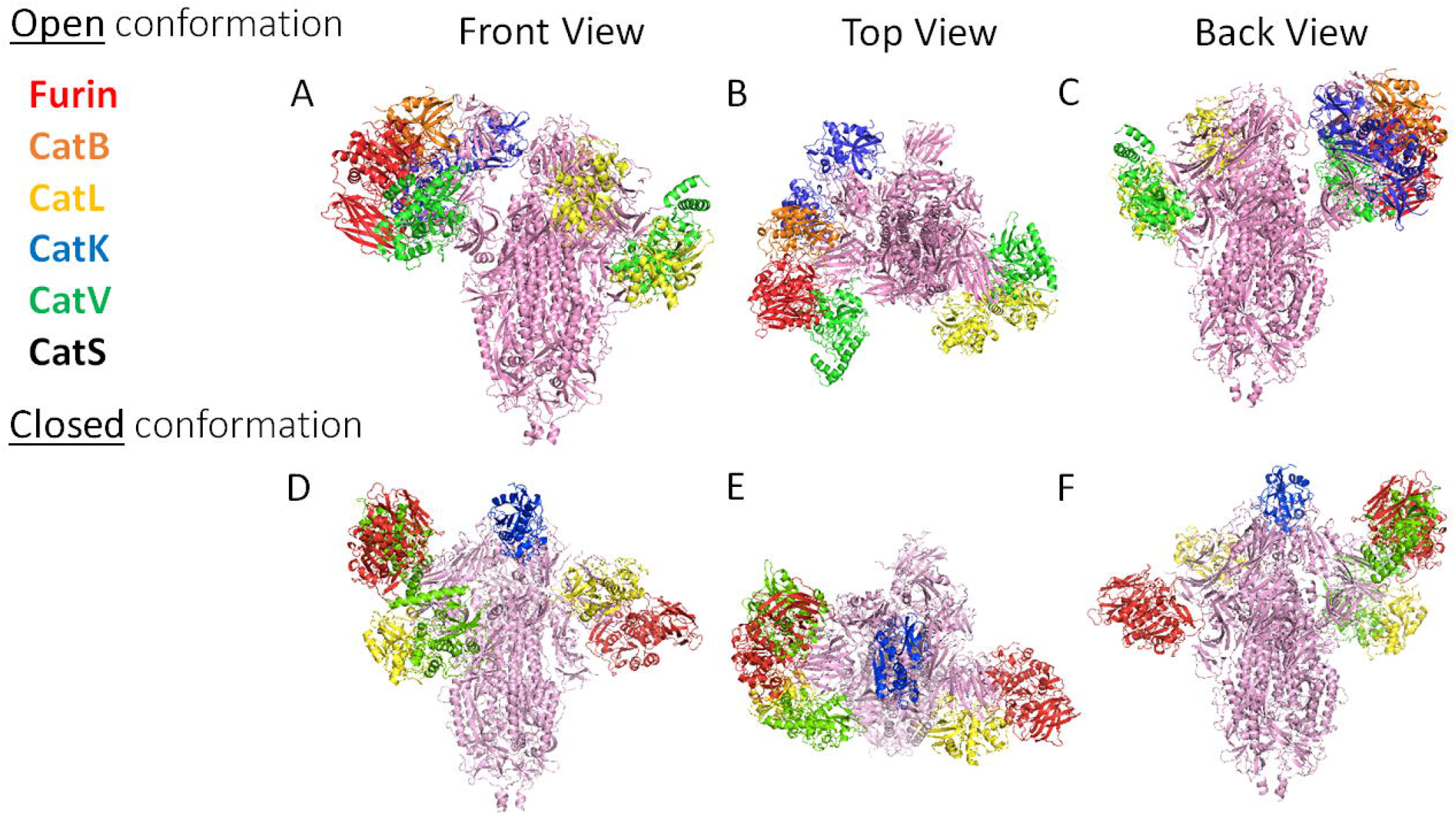
Closest molecular docking sites of each protease on SARS-CoV-2 Spike protein. From the open conformation, molecular docking sites with bound proteases are compiled onto one spike protein trimer and shown from a front (A), top (B), and back (C) view. Furin: 12.2 Å from N81; Cat L: 3.6 Å from V213 and 5.8 Å from A123; Cat B: 10 Å from E340; Cat V: 12.2 Å from R214 and 14.9 Å from A123; Cat K: 3.8 Å from A372 and 11.1 Å from L226. From the closed conformation, docking is shown from a front (D), top (E), and back (F) view. Furin: 14.1 Å from S689 and 14.6 Å from N81; Cat K: 6.2 Å from V367; Cat L: 5.1 Å from R214 and 5.3 Å from A123; Cat V: 11 Å from R214 and 10 Å from N122.

### Putative cleavage of SARS-CoV-2-S protein at S1/S2 and S2’

The spike protein has two key domains, S1 subunit contains angiotensin I converting enzyme 2 (ACE2) receptor binding domain and S2 domain mediates membrane fusion^28^. S1/S2 site is the critical site where cleavage of spike protein by proteases causes a conformational change promoting virus entry to the host cells. The locations of putative maturation sites S1/S2 site 1 and site 2 as well as S2’ on the SARS-CoV-2 spike protein sequence were detailed by Coutard et al.^5^. The regions surrounding these cleavage sites were highlighted areas of analysis. From PACMANS, the rankings are represented by a heatmap with green indicating higher probability of cleavage and red indicating lower. Furin ranked highest at the previously identified furin cleavage sites S1/S2 Site 1 and S2’ (yellow stars), as expected (Figure 2). Cathepsins K, L, S, and V also shared high PACMANS rankings just outside the canonical S1/S2 Site 2, at MSLG/AENS (blue star), suggesting these proteases may also be involved in cleavage that might induce conformational change and membrane fusion. Additional sites such as VTLA/DAGF (pink star) also demonstrated high probabilities of cleavage across multiple cathepsins, with rankings in the top 20 for cathepsins B, K, L, S, and V (Figure 2).

**Figure 2.**
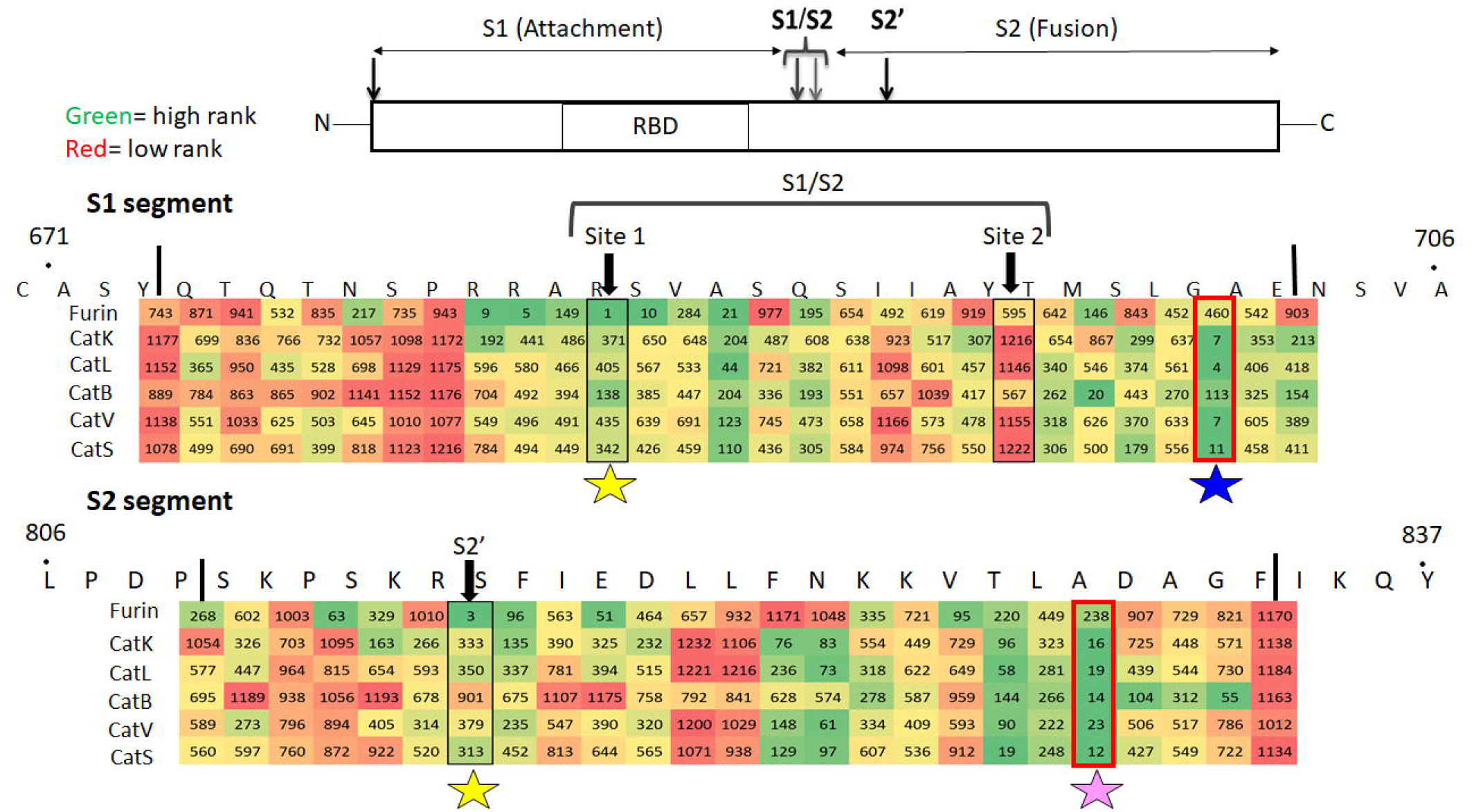
Identification of putative cleavage sites in the S1/S2 region of spike protein based on PACMANS scoring. For each protease, spike protein cleavage sites in the S1 and S2 regions were ranked using PACMANS, and scores are shown with a heat map. Heat scores of ranks are colored from green to red, with green indicating a higher rank (closer to 1) and red indicating lower rank (greater numbers) that are less likely to be susceptible to cleavage by the specific protease. Boxed sequences indicate published cleavage sites of spike protein that promote conformational change for entry to cell. Stars indicate locations where multiple proteases have higher scores of cleavage probability: yellow for furin, blue for the cathepsins K, L, S, and V near S1/S2 site, and pink for cathepsins K, L, S, and V cleaving outside the S2’ site.

### Spike protein sequences susceptible to cleavage by multiple proteases

All four cathepsins, cathepsins K, S, L and V showed high ranks for two of the identified sequences on spike protein, so these were investigated more closely. SVLH/STQD (46-53) in N-terminus domain and MSLG/AENS (697-704) just outside the S2’ site were the two sequences (Figure 3). Three of the cathepsins showed high ranked scores for other sequences. SALE/PLVD (221-228) was scored to be susceptible to cleavage by cathepsins S, V and K in the N-terminus domain (Figure 4A). Cathepsins S, L, and V also showed high ranks for ELLH/APAT (516-523) in the receptor binding domain, and NGLT/VLPP (856-863) in the S2 domain (Figure 4B,C).

**Figure 3.**
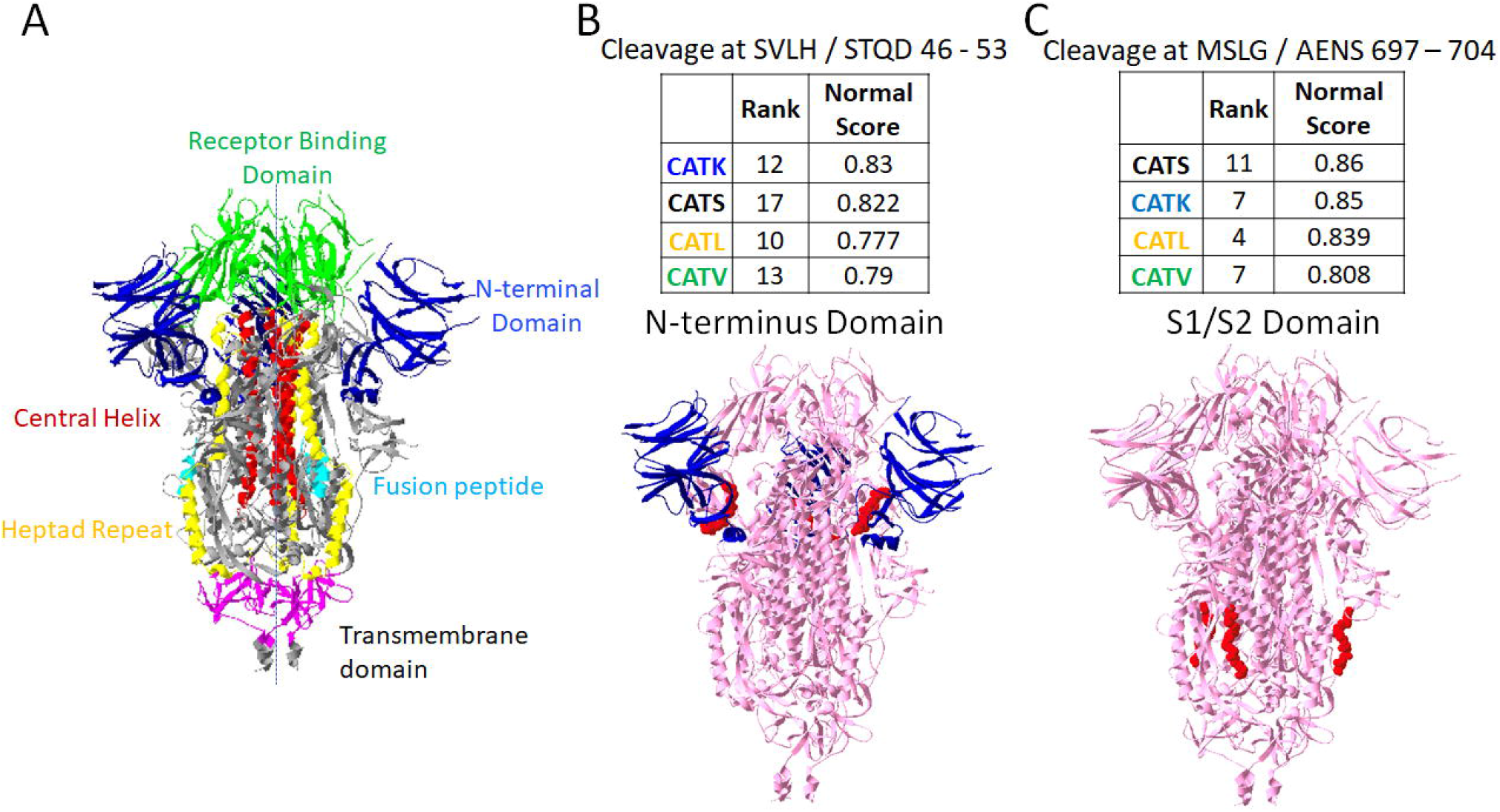
Putative sites susceptible to cleavage by four cathepsins. From PACMANS analysis, there were two amino acid sequences susceptible to cleavage by four cathepsins as indicated by high normalized scores and relatively high rank orders. From the three-dimensional model of spike protein, the domains are color coded, (A) and susceptible sequences are highlighted in red. (B) Cleavage after H49 is in the N-terminus domain (blue) of spike protein, and (C) cleavage after G700 was even more highly ranked and cleaves in the S1/S2 domain.

**Figure 4.**
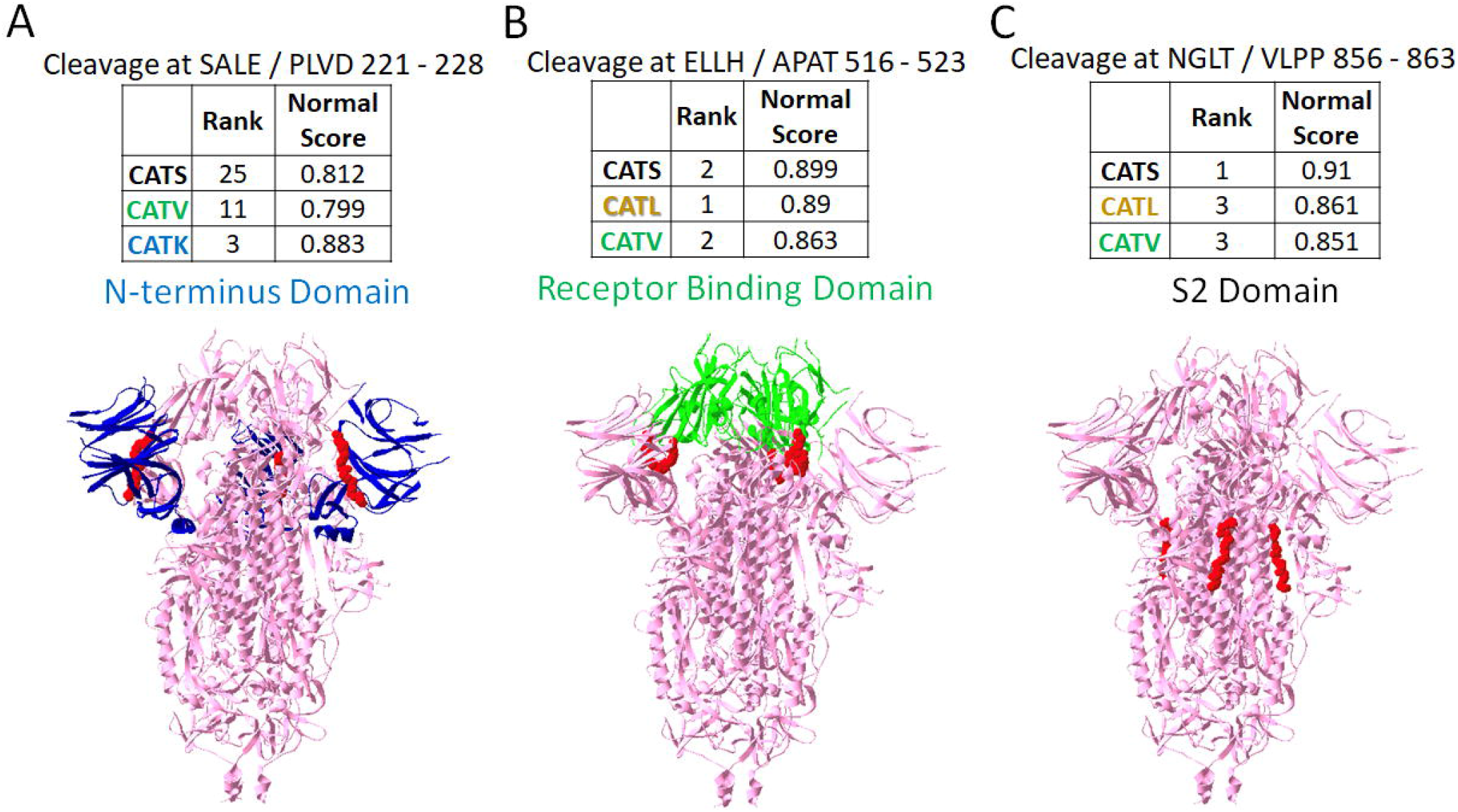
Putative sites susceptible to cleavage by three cathepsins. From the PACMANS analysis, several eight amino acid sequences were susceptible to cleavage by three cathepsins as indicated by high normalized scores and relatively high rank orders. (A) Cleavage after E224 was in the N-terminus domain of spike protein (blue). (B) Cleavage after H519 was even more highly ranked and cleaves in the receptor binding domain (green). (C) Cleavage after T859 was in the S2 domain. Susceptible sequences are highlighted in red.

### Susceptibility for inactivating cleavage of spike protein by multiple cathepsins

With multiple cathepsins having preferred cleavage sites on SARS-CoV-2 spike protein, we proposed the hypothesis that cleavage by cathepsins at multiple sites could be inactivating cleavage events. For example, if two proteases hydrolyzed spike protein at adjacent regions, freeing a part of the protein, then the peptide fragment released could be destabilizing to protein conformation. Using high ranked sequences and molecular docking, cleavage events that would produce fragments were examined. We hypothesize that some of these cleavage events in the N-terminal domain and receptor binding domain might be inactivating, preventing virus binding or cell entry. Using these analyses, fragments to be released from the N-terminal domain were determined and their molecular weights predicted: S50 – V213 (19 kDa fragment), S50 – E224 (21 kDa fragment), S50 – I119 (8 kDa fragment) are in the N-terminal domain and included one of the sites susceptible to cleavage by cathepsins K, L, S, and V (Figure 5A-C). V120 – E224 (12 kDa fragment) is a sequence susceptible to cleavage by cathepsins S, V, and K. Cleavage of these sites and release of these fragments could affect the total protein structure in a destabilizing or unfolding manner that could reduce spike protein’s ability to bind to the ACE2 receptor.

**Figure 5.**
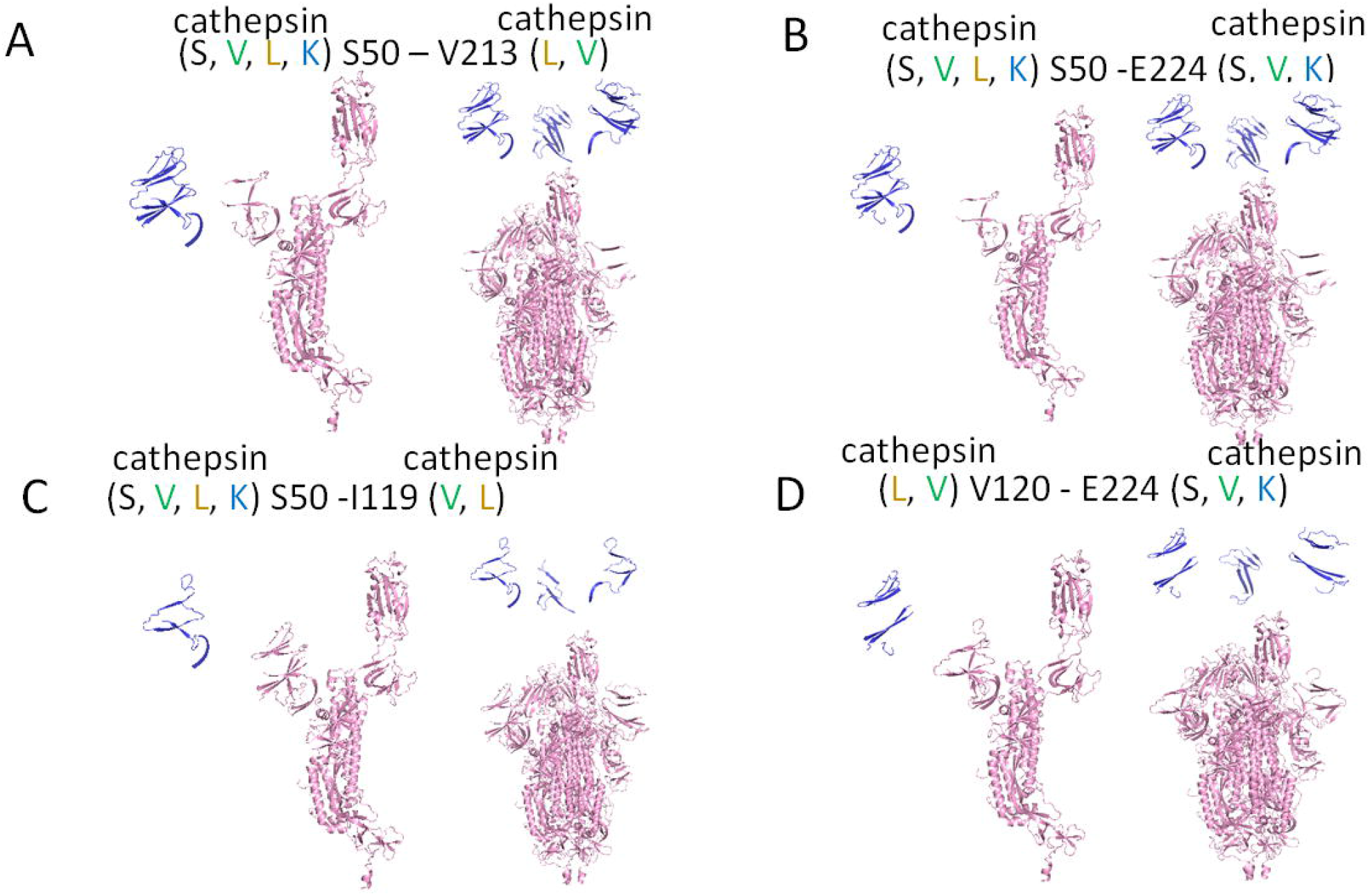
Fragments of spike protein generated due to cleavage at cathepsin putative sites in the N-terminal domain. From PyMol, fragments created from cleavage by multiple cathepsins at two sites on spike protein are shown for (A) S50 through V213, a 19 kDa fragment, (B) S50 through E224, a 21 kDa fragment, (C) S510 – I119, an 8 kDa fragment, and (D) V120 through E224, a 12 kDa fragment. Protomers are shown on the left and spike protein trimers are shown on the right for each cleavage site pair.

For cleavage in the S1/S2 region, again the site susceptible to cleavage by multiple cathepsins were analyzed. G700 – G769 (7 kDa fragment), G700 – V860 (17 kDa fragment), and T768 – V860 (10 kDa fragment) were modeled (Figure 6). Fragments are shown assuming the proteases are cleaving from the protomer of the spike protein, and from the assembled trimer of spike protein. This S1/S2 region is where proteolytic cleavage can be activating, meaning it promotes the conformational change that causes viral membrane fusion with cell membrane and subsequent viral entry. However, if there were multiple cleavage events freeing a fragment from the total spike protein, then the fusion event may not occur when these multiple cathepsins are present.

**Figure 6.**
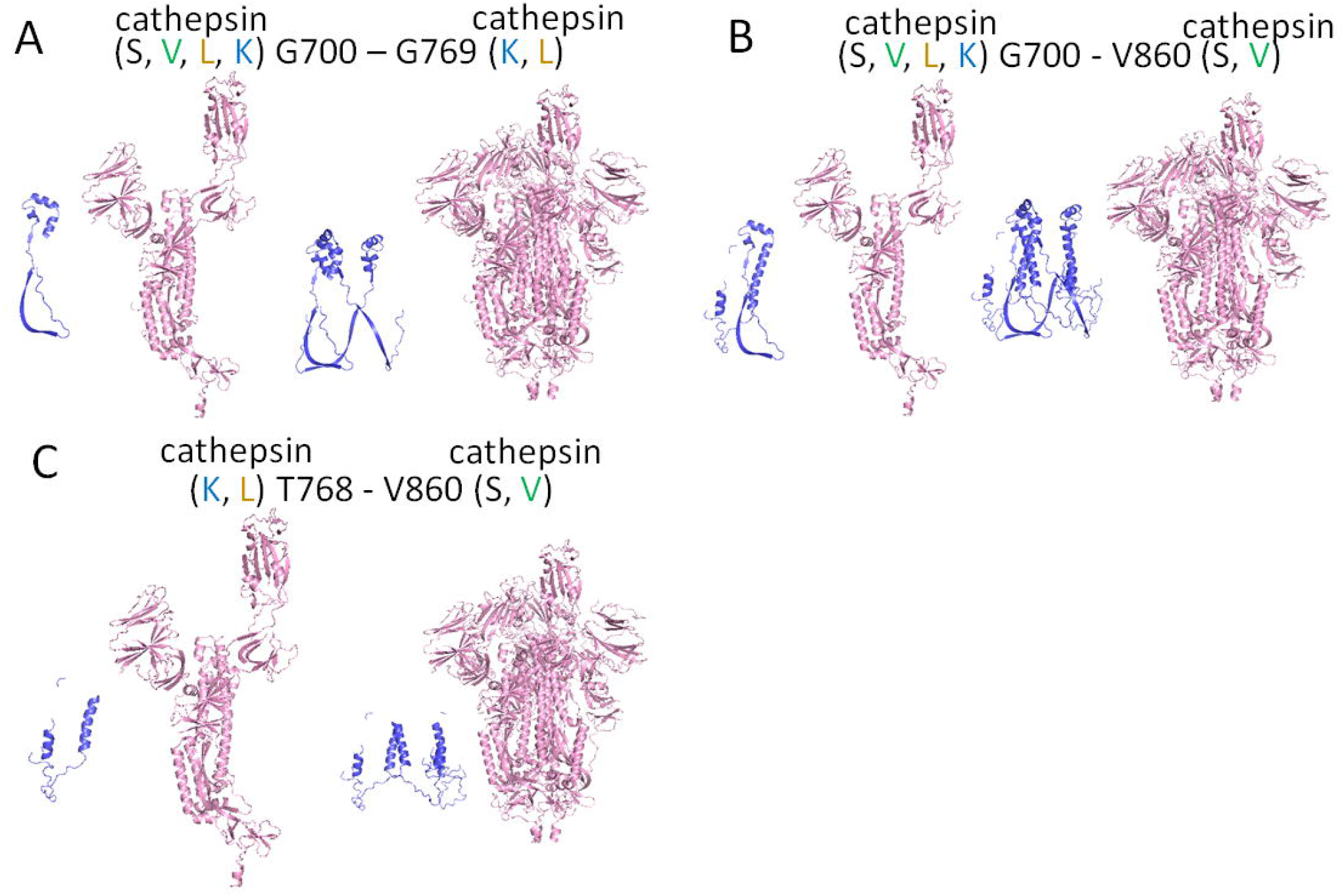
Fragments of spike protein generated due to cleavage in S1/S2 domains. From the docking software, fragments created from cleavage by two or more cathepsins at two sites in the S1/S2 region of spike protein are shown for (A), G700 – G769, 7 kDa fragment (B), T768-V860, a 10 kDa fragment and (C) G700 – V860, a 17 kDa fragment. Fragments are shown from the protomer and from the trimer for each cleavage site pair.

### SARS-CoV-2 spike protein cleavage by individual cathepsins generate unique cleavage fragments

To assess the results from our computational analyses, we conducted experiments testing cleavage of spike protein by cathepsins B, K, L, S, and V. To do this, recombinant spike protein was incubated separately with each cathepsin for increasing periods of time in separate aliquots. Then these samples were probed for immunoblots against SARS-CoV-2 spike protein to visualize cleavage fragments caused by cathepsin hydrolysis. Cathepsins B, K, L, S, and V generated different cleavage fragment patterns from full length spike protein (Figure 7). The cleavage products obtained by the individual cathepsins are unique which corroborate the implications from PACMANS that there are multiple cleavage sites on spike protein where multiple cysteine cathepsins can act, beyond just cathepsins B and L. From the blot, cathepsins B and L actually generated the fewest number of unique immunodetectable cleavage products whereas cathepsins K and V generated a number of uniquely visible cleavage products.

**Figure 7.**
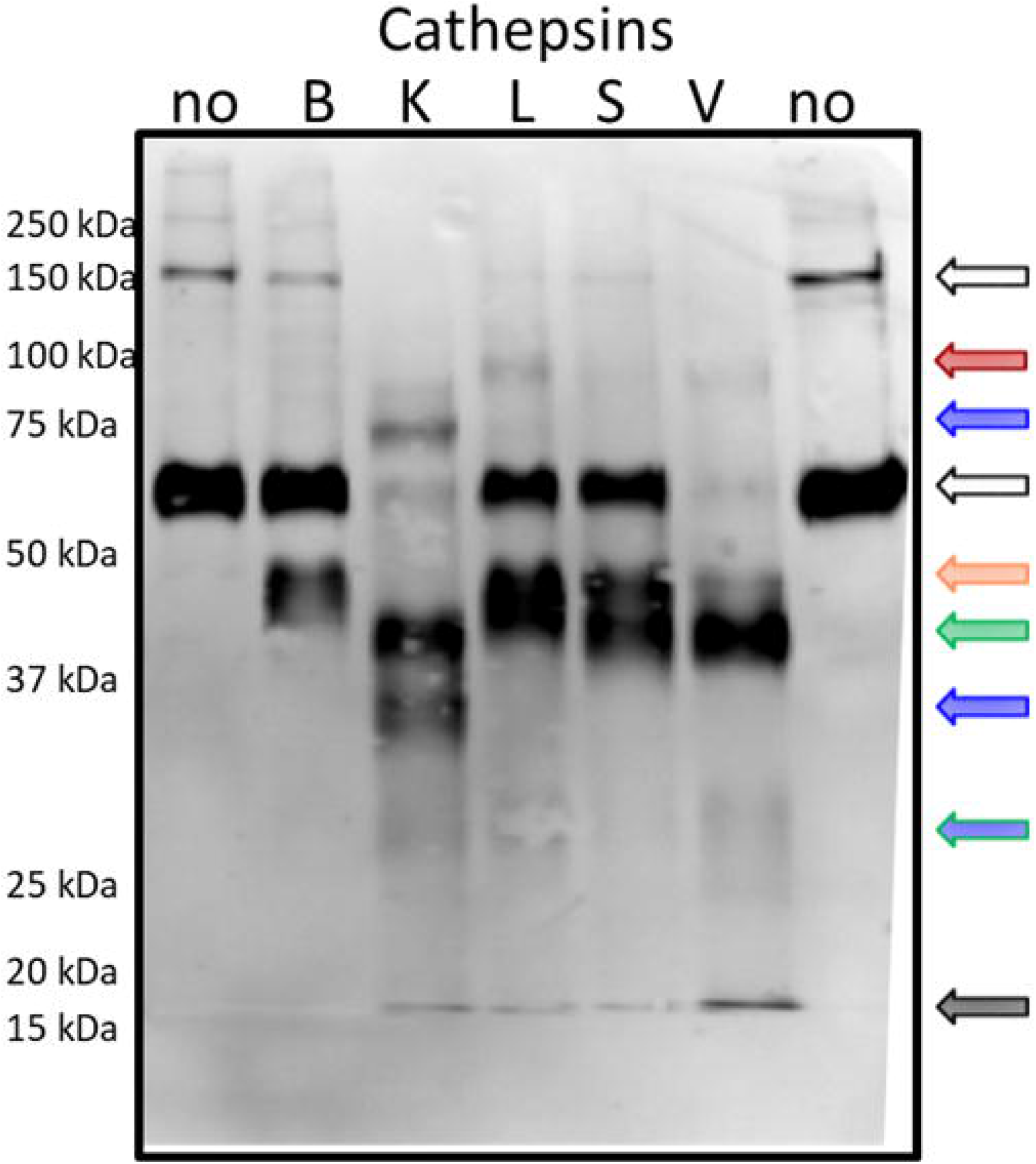
Individual cathepsins cleave SARS-CoV-2 spike protein generating unique cleavage fragments. 5 ng/µl of SARS-CoV-2 spike protein was incubated with 5 pmols of cathepsins B, K, L, S, V or no enzyme for increasing periods of time. Samples were collected and prepared for reducing immunoblot with anti-SARS-CoV-2 spike. Representative blot after 10 minutes of incubation is shown.

The fragment sizes generated reflect some of the predicted cleavage sizes suggested by PACMANS analysis. For example, cathepsin K was suggested to cleave spike protein at G700 and G769 which would generate three fragments: 7 kDa, 56 kDa, 78 kDa (Figure 6); a 78 kDa band appears in the blot. Cathepsin S was suggested to cleave spike protein at both G700 and V860 to release a 17 kDa fragment, but also 46 kDa and 78 kDa fragments; while the 78kDa fragment is not detected, there is a band at 46 kDa and the lower 17 kDa. CatV also has this 46 kDa band as it was suggested to cleave at both of these sites as well. Fragments below 20 kDa in size are present for all of the proteases tested except for cathepsin B. A number of other calculations of band sizes from cleavage fragments can be done, but the validation of specific sequences will need to be done by mass spectrometry to specifically identify peptide fragments that, when re-assembled, recapitulate full length spike protein. However, these computational tools help motivate those further analyses with suggestive hypotheses of multiple cathepsin cleavage events.

## Discussion

From our computational-based PACMANS analysis and computational simulation of protein docking analysis, spike protein is susceptible to cleavage by cathepsins B, K, L, S, V, and furin at multiple sites, which can impact the infectivity and cell entry by SARS-CoV-2. This is the first implication of cathepsins K, S, and V for a role in COVID-19, but we have shown in this computational study that they can potentially cause a maturation cleavage event near the S2’ site that facilitates membrane fusion. A table containing a rank ordered list of sites where cathepsins might cleave spike protein was generated, and we identified two locations that cathepsins K, L, S and V all showed high sequence cleavage preference. As summarized in Table 1 and figures 3 and 4, there are various regions in the spike protein that are accessible and susceptible to cleavage by multiple cathepsins other than S1/S2 domains. N-terminus region and receptor binding domain cleavage sites were identified that could also inactivatingly hydrolyze the spike protein by removing fragments necessary for binding or for destabilizing the spike protein structure. These computational predictions motivated experimental testing of spike protein cleavage by cathepsins B, K, L, S, and V, then immunoblotting confirmed a number of unique fragments of spike protein generated by cathepsin hydrolysis of the full length protein. Taken together, these results have generated a number of hypotheses that can be confirmed experimentally, but also can guide mutational analyses to investigate other potential loss or gain of function mutations in SARS-CoV-2 spike protein using the informed tools described in this study.

Cleavage of the spike protein can occur at the cell surface or in the endolysosomes which puts this cleavage event in two very different microenvironments. At the cell membrane, membrane bound protease, TMPRSS2, or other soluble proteases such as furin have been implicated, and in the acidic microenvironment, cathepsins B and L were shown to be key. While cathepsins play key roles in protein turnover in endosomes and lysosomes, they are also overexpressed and secreted in a number of diseases^11,29^, including those that are pre-existing conditions escalating risk of death due to COVID-19. These conditions could elevate the levels of other cathepsins working in the immediate extracellular spaces near the cell surface where they might facilitate spike protein activation or mitigate its actions by inactivatingly cleaving spike protein. Indeed, cathepsin L has been shown to play a role in protecting from influenza infection through an intracellular mechanism, so there is precedent for proteases acting on both sides^30^. To determine this possibility, we undertook this study of an unbiased search for cathepsin cleavage sites along the spike protein, and then supported those finding with molecular docking analyses and recombinant protein cleavage analyses.

Identification of cathepsin K, L, S, and V cleavage sites in the S1/S2 fusion peptide region suggests their assistance in activating cleavage of spike protein to promote membrane fusion. In addition to the sites discussed, there are two more sequences that are highly susceptible to cleavage by cathepsins K, L V, and S, in which cleavage sequences are located internally on the spike protein in either fusion peptide region (VLTA/DAGF) or heptad repeat region (SALG/KLQD). Cleavage in the fusion protein region of SARS-CoV by cathepsin L, not at the canonical site, still promoted membrane fusion and viral entry^8^ so these locations could later be shown to affect spike protein function as well. As more is learned about the mechanisms of spike protein folding and membrane fusion, then the significance of these protein sequences, and the impact of their proteolysis on either SARS-CoV-2 infectivity or function may become evident.

Cardiac injury has been an important factor associated with mortality of COVID-19^31^. ACE2, the cell receptor that binds spike protein, has previously been shown to be upregulated in cardiomyocytes in patients with cardiovascular disease^32^, and SARS-CoV-2 infection of induced pluripotent stem cell-derived cardiomyocytes has been confirmed^33^. In a separate study, RNA sequencing showed that ACE2 as well as cathepsins B and L where highly expressed in human induced pluripotent stem cell-derived cardiomyocytes, but interestingly, the protease TMPRSS2 was only expressed at low levels^34^. This shows that COVID-19 infection in cardiomyocytes may proceed through cathepsin-mediated pathways more so than TMPRSS2-mediated mechanisms; therefore, confirmation studies need to be completed. It is well reported that protease inhibitors reduce viral infection. FDA approved protease inhibitors used to suppress COVID-19 infection found that ritonavir was most effective at inhibiting SARS-CoV-2 main protease and human TMPRSS2. Cathepsins B and L were also capable of being blocked by indinavir and atazanavir^35^. Again, the other proteases implicated by the results of this study have not been tested but may also be targeted by these inhibitors as well, as cathepsin inhibitors have been notoriously cross-reactive^36–39^.

Based on the computational analysis of this study, sites were identified where proteolysis might be an inactivating cleavage event. These sites were visualized in figures 5 and 6 where adjacent cleavage sites, susceptible to proteolysis by multiple cathepsins, could generate fragments of the spike protein. Then the varying fragments generated by incubation of spike protein with the individual cathepsins, confirmed multiple cleavage events were possible, at least *in vitro*. Once these fragments were removed from spike and the viral capsid surface, then impaired binding to ACE2 receptor or impaired membrane fusion could be sufficient to prevent infection by that virion. Mature, active cathepsins are much smaller than spike protein (∼25 kDa vs. 140 kDa), so while it may be possible that multiple cathepsins could bind to one spike protein, it is more likely that with the distribution of spike proteins along the surface of the virus capsid, that cathepsins would be binding to and hydrolyzing individual spike proteins iteratively, but subsequent hydrolytic events by multiple proteases could lead to the results predicted in figures 5 and 6 and then shown in figure 7. This would also be dependent on concentrations of proteases present in the extracellular environment, in a tissue-specific manner, where virions are trying to attach to cell surfaces. Patients with cardiovascular disease showed dysregulated inflammation^40^, increased permeability^41^, upregulated protease^42^ and epithelial dysfunction^43^. Severe inflammation, such as the cytokine storm triggered by COVID-19^44^, can also cause an influx of cysteine cathepsins produced by inflammatory macrophages, alveolar macrophages, endothelial cells, and smooth muscle cells, all of which produce cathepsins in response to cytokines such as TNFα, IFNγ, IL-1β^19,45–47^.

Though we find this study to be useful, there are limitations to be considered. The results presented identify hypothetical protease cleavage sites on SARS-CoV-2-S, and spike protein hydrolysis by cathepsins K, S, and V was verified with recombinant *in vitro* studies. However, confirmation of the identity of these sequences to confirm they match with PACMANS predictions must be completed with mass spectrometry or site-directed mutagenesis for confirmation. The benefits of the computational approaches applied here are that a process has been presented of PACMANS followed by molecular docking to generate hypotheses for investigating other proteases, even beyond those included in this study. Upregulated plasmin, thrombin, and other serine proteases may be present at the site as well, implicated in the hypercoagulable state induced by SARS-CoV-2 infection^48^. Another example involves the D614G SARS-CoV-2 spike mutant that has been shown to be a more infectious virus with an enhanced susceptibility of spike protein to furin cleavage, while also altering the dynamics of the spike protein conformation^49^. This site was not identified in the unbiased approach used in the studies presented here, but if other mutations are identified, they can be input into PACMANS to determine their putative result on activating cleavage. Another study identified a cleavage site on spike protein from SARS-CoV, the virus that caused the SARS outbreak in 2009, at position T678^8^, but that site also was not highly ranked in the unbiased approaches used here. Despite these sites not appearing in the analysis, we are confident that the PACMANS analysis identifies cleavage sites as the furin site was the top ranked site, and additional cleavage sites that had been validated by other experimental approaches were identified from this analysis.

The extension from these analyses would be the further discovery of the role multiple cathepsins interacting, binding, and cleaving spike protein in activating or inactivating ways that can be taken in the aggregate to predict the responses in cells and tissues when exposed to SARS-CoV-2. We also have identified interactions of cathepsins with each other that remove them from the system, as they work in a proteolytic network and affect substrate degradation^50^. Together, these systems considerations of multiple cathepsins working with or against other proteases in the presence of spike protein during SARS-CoV-2 infection may yield insight into optimizing pharmaceutical targeting to mitigate the effects.

## Materials and Methods

### PACMANS Analysis to identify putative cleavage sites

Protease-Ase Cleavage from MEROPS Analyzed Specificities (PACMANS) algorithm developed by Ferrall-Fairbanks et al. was utilized to rank potential cleavage sequences^25^. Protease specificity matrixes were obtained from the MEROPS peptidase database. PACMANS ranks each eight amino acid sequence in the substrate by its likelihood of hydrolysis by the selected protease.

### 3D Molecular Docking Simulation

The possible binding of SARS-CoV-2-S protein (PDB ID: 6VYB) and furin (4Z2A), cathepsins L (5F02), cathepsin S (4P6E), cathepsin K (5TUN), cathepsin B (2IPP), and cathepsin V (3H6S) were simulated using Clus Pro 2.0, in which 3D protein crystal structures from protein data bank were automatically stripped of their water molecules and additional ligands before beginning docking simulations^51,52^. Each simulation yielded 15-30 possible docking models using a balanced binding interaction (electrostatic, hydrophobic, and Van der Waals) model. Top 10 simulation models were viewed in PyMOL followed by highlighting the active sites of catalytic triad of each protease. Among those, only the models that catalytic triad facing the SARS-CoV-2 S protein were further analyzed for possible cleavage. The distance was measured in Angstroms between the closest residue on the S protein and the catalytic triad. Distances of less than 10 Angstroms were considered of most interest.

### SARS-CoV-2-S protein domain color coding

The N terminal domain, receptor binding domain, fusion peptide, heptad repeat 1, central helix, and connector domains of each subunit of the SARS-CoV-2-S protein were identified in SwissPBD by comparison of the SARS-CoV-2 PDB sequence and the domain sequences outlined by Wrapp et al.^53^ Each domain was colored in the SwissPBD model to visualize potential cleavage site locations in relation to these domains.

### Cathepsin hydrolysis of SARS-CoV-2 spike protein and immunoblotting

Five pmol of recombinant human cathepsins B, K, L, S, and V (Enzo) was incubated with 5 ng/µl of SARS-CoV-2 (2019-nCoV) Spike S1+S2 ECD-His Recombinant Protein (Sino Biological #40589-V08B1) in phosphate buffer (pH 6.0), 2 mM DTT, 1 mM EDTA for at 37°C for defined time periods. Reducing SDS-PAGE loading buffer was added and then heated at 95°C to terminate experiments. After SDS-PAGE, protein was transferred to a nitrocellulose membrane (Bio-Rad) and probed with SARS-CoV/SARS-CoV-2 Spike Protein S2 Monoclonal Antibody (1A9) (ThermoFisher # MA5-35946) and donkey anti-mouse secondary antibodies tagged with an infrared fluorophore (Rockland Immunochemicals). Membranes were imaged with a LI-COR Odyssey scanner.

## Acknowledgments

This work was supported by the National Science Foundation through Science and Technology Center Emergent Behaviors of Integrated Cellular Systems (EBICS) Grant CBET-0939511 (M.O.P).

